# Canonical Wnt signalling from the Area Opaca induces and maintains the Marginal Zone in pre-primitive-streak stage chick embryos

**DOI:** 10.1101/2024.08.24.609497

**Authors:** Yara Fadaili, Hui-Chun Lu, Hyung Chul Lee, Amra Ryazapova, Claudio D. Stern

## Abstract

In chick embryos prior to primitive streak formation, the outermost extraembryonic region, known as the area opaca (AO), was generally thought to act only by providing nutrients and mechanical support to the embryo. Just internal to the AO is a ring of epiblast called the marginal zone (MZ), separating the former from the inner, area pellucida epiblast. The MZ does not contribute cells to any part of the embryo but is involved in determining the position of primitive streak formation from the inner epiblast. Recently it was discovered that the AO can induce a MZ from area pellucida epiblast. Here we explore the nature of this inductive signal. We find that Wnt8c is highly expressed in the AO, whereas canonical Wnt pathway targets are enriched in the MZ, along with strong nuclear β-catenin localization. Using isolation and recombination experiments combined with gain- and loss-of-function by exogenous chemical modulators of the pathway, we reveal that Wnt signalling is essential for induction and maintenance of the MZ, as well as sufficient to induce MZ properties in area pellucida epiblast. We propose that canonical Wnt signalling is responsible for induction of the marginal zone by the area opaca.

## Introduction

The chick embryo at stage X, when the egg is laid, is a flattened disc-shaped blastoderm with a continuous epiblast layer that is divided into three concentric regions (Fig. 1A). The central region, known as the area pellucida (AP), contains all cells that will contribute to the embryo (Eyal-Giladi and Kochav, 1976, Hatada and Stern, 1994, Lee et al., 2020). A ring of epiblast immediately surrounding the AP (delimited from it posteriorly by Koller’s sickle) is called the marginal zone (MZ). Its cells do not contribute to any embryonic tissues, but it is involved in the establishment and maintenance of embryonic polarity, specifically by determining the location at the posterior edge of the AP, where primitive streak formation will begin (Eyal-Giladi and Khaner, 1989, Eyal-Giladi and Kochav, 1976, Bachvarova et al., 1998, Lee et al., 2020, Torlopp et al., 2014, Lee et al., 2024b). The outermost region, called the area opaca (AO), which is also extraembryonic, has generally been considered to provide nutrients and mechanical support for blastoderm expansion but otherwise to have no instructive role in morphogenesis of the embryo (Bellairs et al., 1967, Downie, 1976, Lee et al., 2022a, Lee et al., 2024a).

**Fig. 1.**
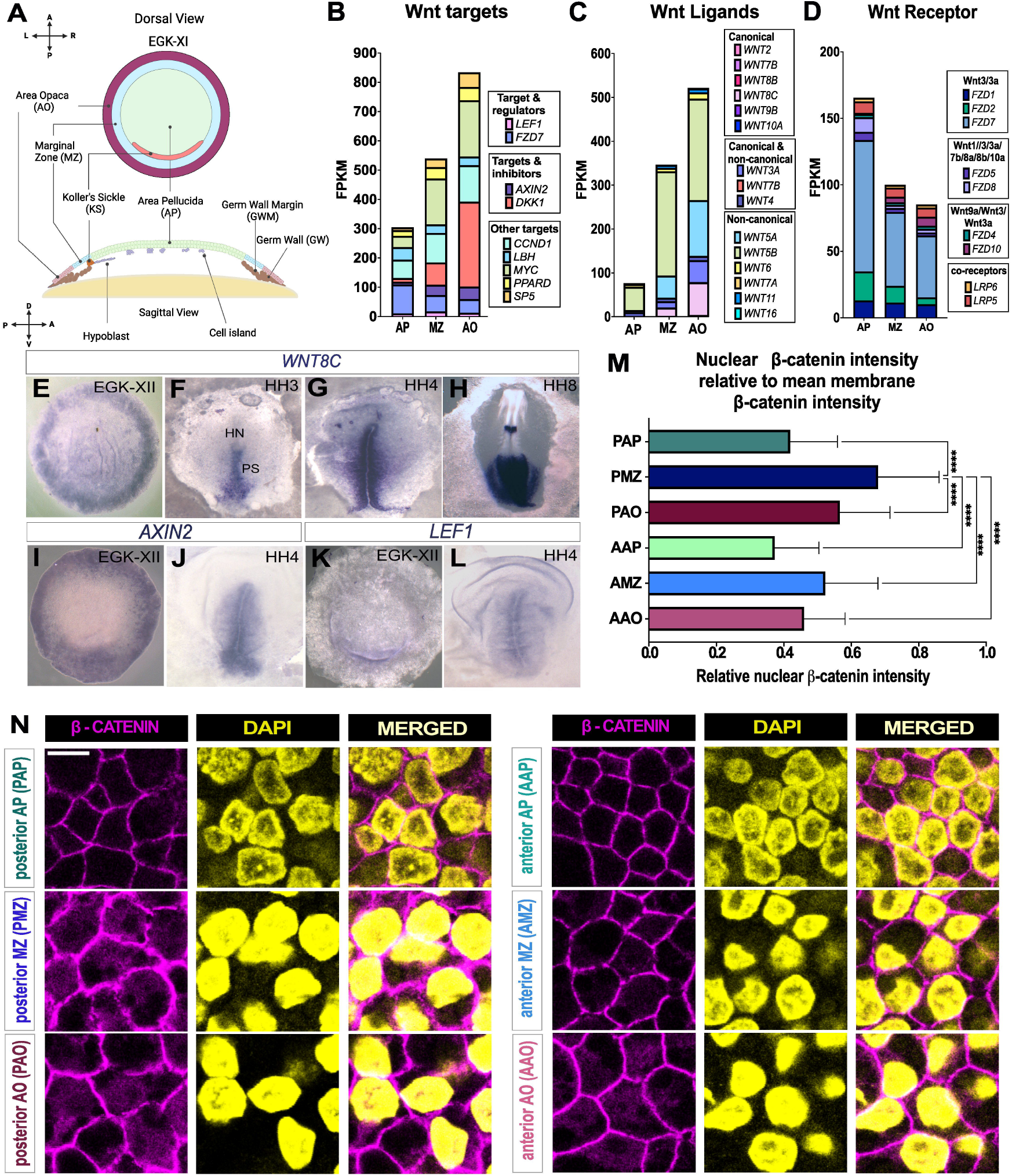
Spatial and regional expression of canonical Wnt ligands, targets, and effectors. **A**. Schematic diagram of the different regions of a stage EGK-XI embryo. **B-D**. Stacked bar chart of RNA-sequencing data analysis of different regions collected from stage EGK-XII embryos (Lee et al., 2020). Whole AO, whole MZ and whole AP were analysed to compare FPKM values of Wnt related molecules, grouped into 3 categories: Wnt targets (B), Wnt ligands (C) and Wnt receptors (D). **E-L**. In situ hybridisation of canonical Wnt ligand WNT8C at different stages EGK-XII (E), HH3 (F), HH4 (G) and HH8 (H). **I-L**. In situ hybridisation of Wnt targets AXIN2 (I, J) and LEF-1 (K, L) in embryos at stages EGK-XI (I, K) and HH4 (K, L). **M-N**. Regional differences in β-catenin localization in early pre-gastrulation chicken embryo. **M**. Bar graph representing nuclear β-catenin pixel intensity (obtained from nuclear segmentation of DAPI) relative to the mean membrane intensity (obtained by membrane β-catenin segmentation) (n=3) (Kruskal-Wallis t-test; P-value < 0.0001=****) between all regions compared to the PMZ. **N**. β-catenin staining 3-slice maximum projection; scale bar= 10 μm. AAO: anterior area opaca, PAO: posterior area opaca, AMZ: anterior marginal zone, PMZ: posterior marginal zone, AAP: anterior area pellucida, PAP: posterior area pellucida. AAO: anterior area opaca, PAO: posterior area opaca, AMZ: anterior marginal zone, PMZ: posterior marginal zone, AAP: anterior area pellucida, PAP: posterior area pellucida.

A new role for the opaca has recently been uncovered. When the MZ is removed and the AO made to surround the central AP directly, a new MZ is induced, expressing specific markers such as *ASTL* (Lee et al., 2022b). This finding suggests that the area opaca plays a role in establishment of the marginal zone and thereby positioning of the primitive streak. The question remains as to the nature of the signalling molecules emitted by the AO that are responsible for the induction of MZ character.

Here we address this question. First, we survey expression of secreted factors and their corresponding receptors in these three different regions of the early embryo using by mapping CellChatDB (Jin et al., 2021) annotations to the RNAseq data and by in situ hybridisation. This initial survey points to canonical Wnt as a possible candidate. Indeed, we find that targets of this pathway are highly expressed in the MZ, as is nuclear localisation of β-catenin, indicating that this region receives strong Wnt signals in the normal embryo. Next, we explored the role of canonical Wnt in the AO more directly by loss-of-function using the Tankyrase inhibitor IWR-1, and gain-of-function with the GSK-3 inhibitor BIO. Wnt inhibition hindered the ability of the AO to induce a new MZ from AP epiblast, whereas BIO increased the inductive ability of the AO. When an isolated AP is treated with BIO, it loses AP markers and gains expression of MZ markers, suggesting that Wnt is sufficient to transform the entire AP into a MZ. Moreover, normal, intact stage X embryos, which possess an established MZ, lose MZ markers when treated with IWR-1, suggesting that continued Wnt signalling is required to maintain MZ after its establishment in normal development. We therefore propose that canonical Wnt signalling is the MZ inducing signal from the AO. This is reminiscent of the role of canonical Wnt signalling in dorsoventral patterning and for establishing the Nieuwkoop centre in Xenopus embryos (Guger and Gumbiner, 1995, Funayama et al., 1995, He et al., 1995, Yost et al., 1996, Yost et al., 1998, Schneider et al., 1996).

## Results

### High Wnt signalling activity in the area opaca and marginal zone of early chick embryo

To identify candidate signalling pathways that are active in the area opaca and can be received by the responding area pellucida, we started by exploring which ligands are expressed in the area opaca (as the signalling region) that have corresponding receptors expressed in either the area pellucida (which is competent to respond to signals from the area opaca) or the marginal zone (which has presumably been induced by these signals earlier). We used the CellChatDB ligand-receptor annotations (http://www.cellchat.org/cellchatdb/) (Jin et al., 2021) to map gene expression data in fragments per kilobase of transcript per million mapped reads (FPKM) obtained from RNA-sequencing data (Lee et al., 2020) for stage (Eyal-Giladi and Kochav, 1976) (EGK) XII embryos, for the whole area opaca (wAO), whole area pellucida (wAP) and whole marginal zone (wMZ). The results are presented in Supplementary Table S1. Some prominent candidates include Wnt ligands (especially WNT3A, 5A, 5B and 8C) and their Fz receptors, Apela/Aplnr, BMPs and Nodal. The canonical Wnt co-receptors LRP5/6 are also expressed in the responding tissue (Supplementary Table S1).

To verify this prediction, we explored the expression of individual Wnt targets (Fig. 1B), ligands (Fig. 1C) and receptors (Fig. 1D). The average FPKM values of all canonical Wnt targets is 2.7-fold higher in the area opaca and 1.8-fold higher in the marginal zone compared to the area pellucida (Fig. 1B). The FPKM value of Wnt ligands is 1.5-fold higher in the area opaca than in the marginal zone and 6.8-fold higher than in the area pellucida (Fig. 1C). Wnt receptors, on the other hand, show a complementary expression profile, with the combined FPKM values of the receptors in the area pellucida being 1.6 fold higher than in area opaca and 1.2-fold higher than in the marginal zone than in the area opaca (Fig. 1D).

Among the canonical Wnt ligands, *WNT8C* is expressed in the area opaca at 4-fold higher level than in the marginal zone, and no significant expression is detected in the area pellucida (Fig. 1C). This expression pattern was confirmed by *WNT8C* in situ hybridisation of pre-primitive-streak stage embryos (Fig. 1E) and agrees with earlier findings (Hume and Dodd, 1993, Skromne and Stern, 2002). Similarly, the Wnt target and modulator *AXIN2* reveals overlapping expression with *WNT8C. AXIN2* has higher expression in the AO and MZ, but no expression is detected in the AP (Fig. 1I). The expression of another Wnt target, *LEF-1, is* confined to the posterior MZ in EGK-XI embryos (Fig. 1K). Together, these expression patterns suggest that canonical Wnt signals, probably delivered by WNT8C, are produced mainly by the extraembryonic AO and received by the MZ. The high expression of Wnt receptors but not Wnt targets in the area pellucida is consistent with this region being competent to respond to Wnt signals, but that at this stage they are not acting significantly beyond the MZ. We also observed that once the primitive streak forms at stage HH-2 (Hamburger and Hamilton, 1951), *WNT8C* (Fig. 1 F-H), *AXIN2* (Fig. 1J) and *LEF-1* (Fig. 1L) expression are all cleared from the AO and MZ and instead become expressed in the primitive streak itself, suggesting a different role of Wnt signalling at this later stage.

To confirm these findings, we performed immunostaining for the canonical Wnt effector β-catenin. We quantified nuclear β-catenin staining intensity relative to the membrane intensity in segmented images of 6 different regions: posterior-AO, -MZ and -AP, and anterior-AO, -MZ and -AP, respectively. We observed that the highest nuclear intensity of β-catenin is seen in the posterior side of the embryo, and that the AO and MZ (both anterior and posterior sides) show higher nuclear intensity of β-catenin than the AP. The posterior MZ displays the highest nuclear intensity of β-catenin, compared to all other embryonic and extraembryonic regions (P-value <0.0001; Kruskal Wallis test) (Roeser et al., 1999) (Fig. 1 M-N).

Together, these results show that canonical Wnt signalling activity in pre-primitive streak stage embryos is concentrated in extra-embryonic regions, as suggested by previous studies (Lee et al., 2020, Roeser et al., 1999, Schmidt et al., 2004, Skromne and Stern, 2002). The expression profiles of Wnt ligands, receptors and targets implicates Wnt signalling as a good candidate for the recently proposed signal from the area opaca that can induce MZ from responding area pellucida cells (Lee et al., 2022b).

### Canonical Wnt signalling from the area opaca induces marginal zone identity in area pellucida epiblast

The above findings suggest that Wnt signalling from the area opaca may be responsible for the induction of marginal zone from area pellucida epiblast. To test this possibility, we performed ablation and recombination experiments combined with modulation of Wnt signalling. Wnt activity can be inhibited with the Wnt antagonist IWR-1 (25 μM), which inhibits Tankyrase (Martins-Neves et al., 2018, Narwal et al., 2012) or stimulated with the Wnt agonist BIO (10 μM), which inhibits GSK-3 (Meijer et al., 2003) (Supplementary Fig.S1 A). First, we validated the efficiency of these treatments. We incubated EGK-XI embryos with the Wnt modulators for six hours and the expression of the Wnt target *AXIN2* was examined. Treatment with IWR-1 led to significant reduction in *AXIN2* expression in 7/8 embryos, a significant effect compared to embryos treated in vehicle alone (0.2% DMSO; 9/9 with normal expression) (Fisher’s exact test p = 0.0004) (Supplementary Fig. S1 B-E). Conversely, treatment with BIO caused a significant increase in *AXIN2* expression (7/9 embryos) (Fisher’s exact test p = 0.0023) (Supplementary Fig. S1 B-E). The effect of the pharmacological agents was also evaluated by immunostaining for β-catenin. IWR-1 treated embryos showed significantly lower nuclear intensity of β-catenin compared to controls in the posterior AO (Mann-Whitney U test P-value <0.0001; n=2) (Supplementary Fig. S1 F, G, I), while embryos treated with BIO showed significantly higher nuclear intensity of β-catenin relative to controls (Supplementary Fig. S1 F, H, I) (Mann-Whitney U test P-value <0.0001; n=2). These results confirm that BIO and IWR-1 at these concentrations are effective as modulators of Wnt activity.

With these tools, we assessed whether canonical Wnt signalling is required for the AO to induce MZ in AP epiblast (Lee et al., 2022b). The entire circumference of the MZ was excised from stage EGK-X embryos, and the AO (with its posterior portion removed) juxtaposed onto the AP to surround it completely, to generate AO-AP conjugates (Fig 2A), as previously described (Lee et al., 2022b). To check that the MZ had been completely removed, we performed situ hybridisation for the marginal zone marker *ASTL* in EGK-XI embryos immediately after ablation: indeed, no expression remained (Fig. 2B). Induction of a new MZ was assessed after incubation of the AO-AP conjugates 7 hours, by in situ hybridisation for the MZ markers *ASTL* and *GJB6*. We asked whether Wnt activity is necessary for MZ induction by the AO. In control (0.2% DMSO) incubated AO-AP conjugates, a new MZ was induced, marked by expression of *ASTL* (7/12 conjugates) (Fig. 2 C, I) and *GJB6* (7/8 conjugates) (Fig 2. F, I). However, some AP-AO conjugates failed to induce a marginal zone with *ASTL* expression (5/12). When AO-AP conjugates were incubated in Wnt antagonist IWR-1, the induction of MZ by the AO was severely impaired: 6/8 conjugates lacked *ASTL* expression (Fig. 2 D, I) and 3/6 had no *GJB6* expression (Fig. 2 G, J). These results suggest that Wnt signalling is required for AO to induce a new MZ in the AP.

**Fig. 2.**
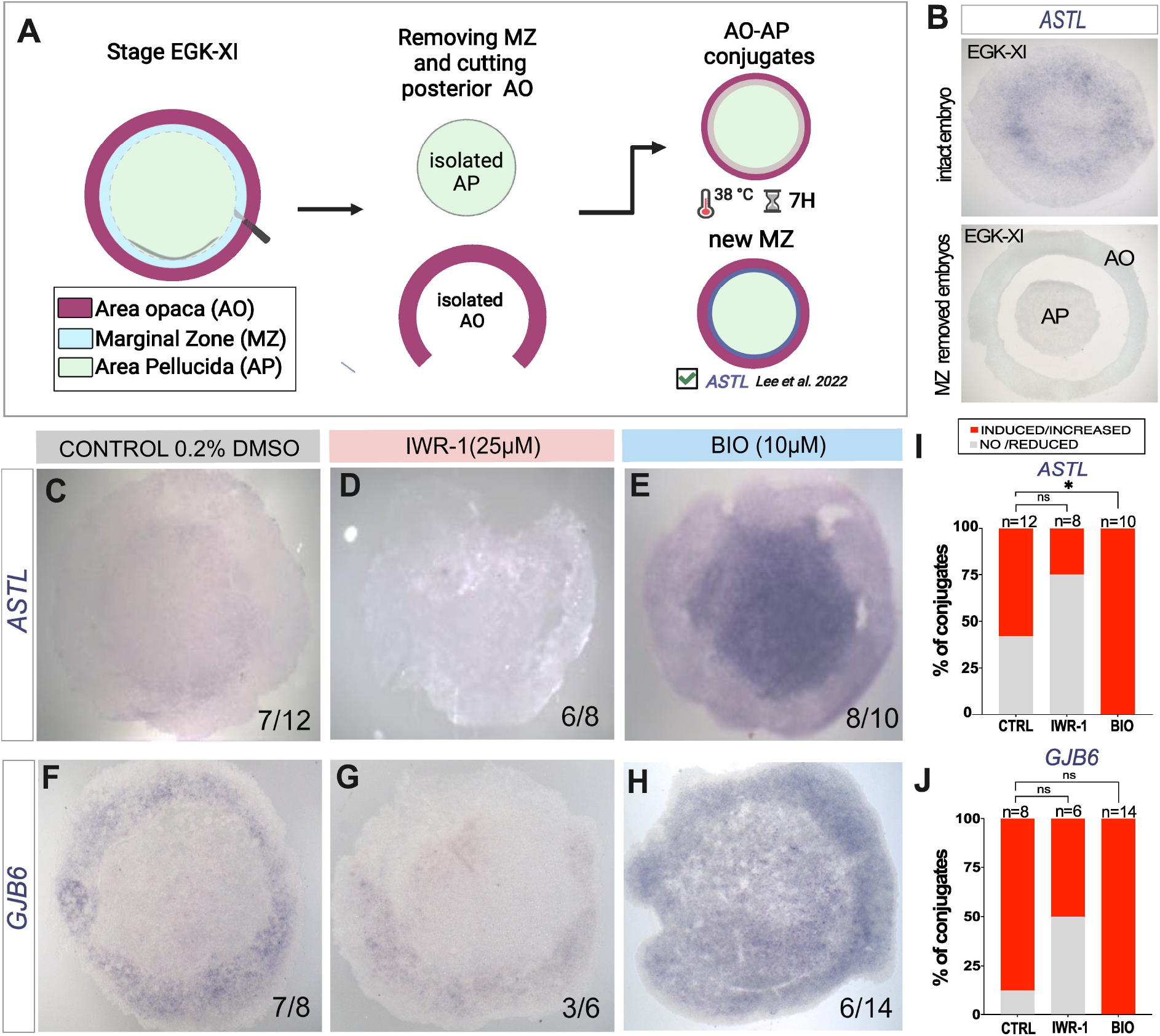
Specification of the MZ and Wnt signalling. **A**. Schematic diagram showing experimental design and possible results when removing the MZ and stitching the extraembryonic AO onto the embryonic AP. **B**. Test for accuracy of ablation: In situ hybridisation of MZ marker *ASTL* in intact and MZ ablated (AO+AO explants) EGK-XI embryos. **C-H**. In situ hybridisation of MZ markers ASTL (C-E) and GJB6 (F-H) in control AO-AP conjugates (0.2% DMSO, C, F), and conjugates treated with IWR-1 (25μM, D, G) or BIO (10μM, E, H) after 7-hours’ incubation. **I-J**. Stacked bar graph representing percentage of AP-AO conjugates with induced or not induced ASTL (I) and GJB6 (L). P-values were determined by Fisher’s test: CTRL vs. IWR (p-value 0.1968; ns), CTRL vs. BIO (p-value 0.0396; *), CTRL vs. IWR-1 (p-value 0.0699; ns), CTRL vs BIO (p-value >0.999). ns: not significant, CTRL: control.

Next, we explored the effect of increased Wnt activity on this inductive interaction by treating AO-AP conjugates with BIO. This resulted in induction of a broader region of marginal zone than in controls as revealed by *ASTL* (8/10 conjugates) (Fig. 2 C, E, I), and *GJB6* expression (6/14) (Fig. 2 F, H, J).

### Wnt signalling is sufficient to induce the marginal zone

Although the results presented so far implicate Wnt in MZ induction, they do not rule out the possibility of another signal within the AO cooperating with Wnt in this process. To investigate if Wnt activity alone is sufficient to induce marginal zone properties, we removed both the MZ and the AO and incubated the isolated AP in BIO for 7 hours before assessing the expression of MZ markers *ASTL* and *GJB6* and of the AP marker *GJA1* (Fig. 3A) by in situ hybridisation. *ASTL* expression was induced ectopically and very broadly in the AP after Wnt stimulation by BIO in 8/9 AP explants (Fig. 3 C, D), whereas control (0.2% DMSO incubated) AP explants either showed no (5/8) or greatly reduced (3/8) expression (Fig. 3 B, D). Moreover, expression of the AP specific gap junction component *GJA1* was inhibited in the BIO treated explants (4/7) (Fig. 4 I, J) while the MZ specific gap junction *GJB6* was upregulated (3/4) (Fig. 3 F, G). In contrast, control (0.2% DMSO) AP explants expressed *GJA1* (11/11; Fig. 3 H, J) but not *GJB6* (4/4; Fig. 3 E, G). These results suggest that Wnt is indeed sufficient to induce MZ identity in area pellucida epiblast, even in the absence of the AO.

**Fig. 3.**
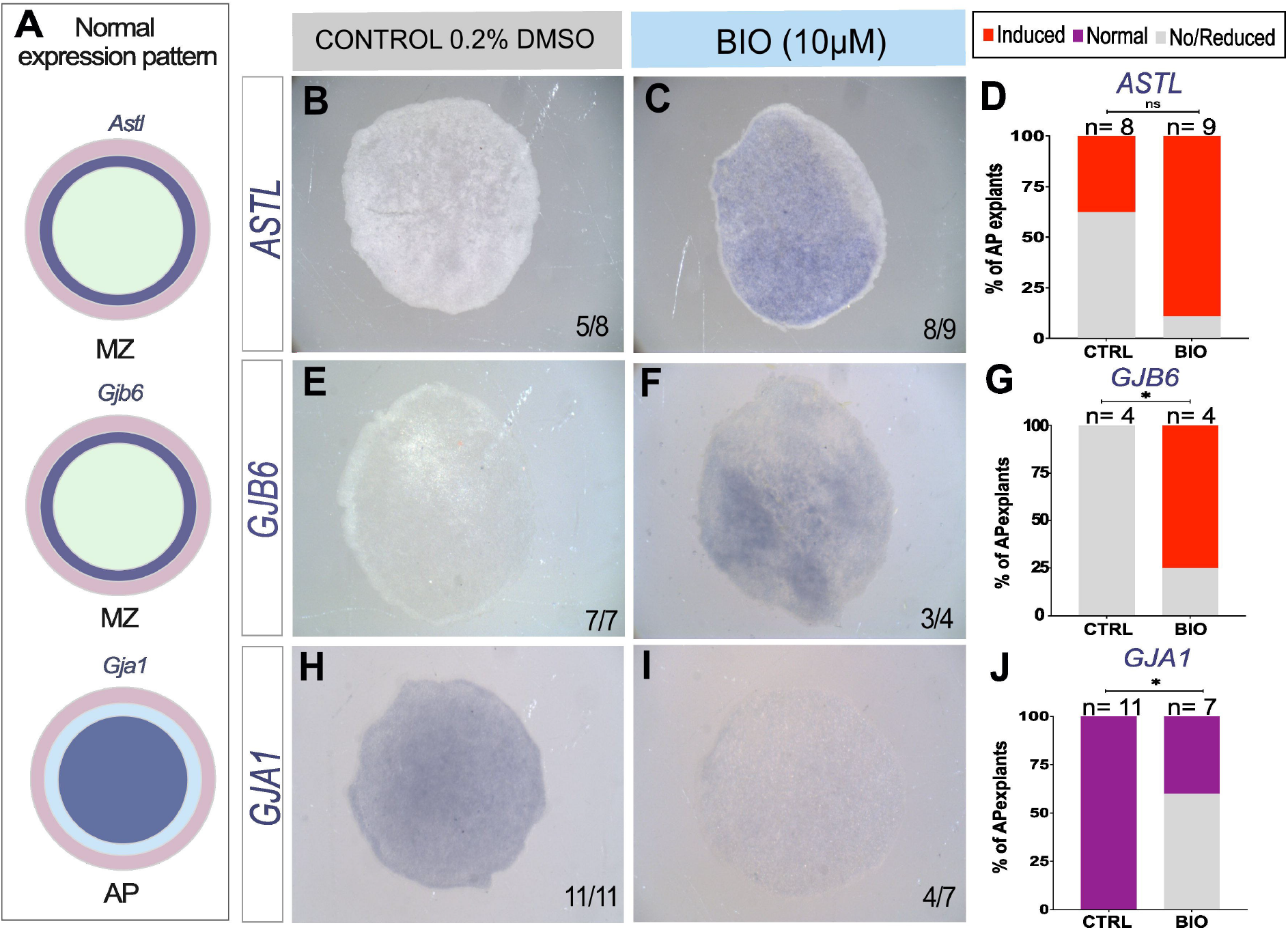
Wnt stimulation induces MZ in AP explants. **A**. Summary of the normal expression domains of ASTL, GJB6 and GJA1. **B-J**. In situ hybridisation in control (0.2% DMSO; B, E, H) and Wnt stimulated BIO-treated (10μM, C, F, I) AP explants after 7 hours’ culture; ASTL (B, C) GJB6 (E, F) and GJA1 (H, I). **D, G, J**. stacked bar graphs showing the percentage of AP explants with induced expression or no expression ASTL (D), GJB6 (G) and Gja1 (J). P-values determined by Fisher’s test: ASTL CTRL vs BIO (p-value 0.3168); ns, GJB6 CTRL vs BIO (p-value 0.0242; *) and GJA1 CTRL vs BIO (p-value 0.0114; *)

**Fig. 4.**
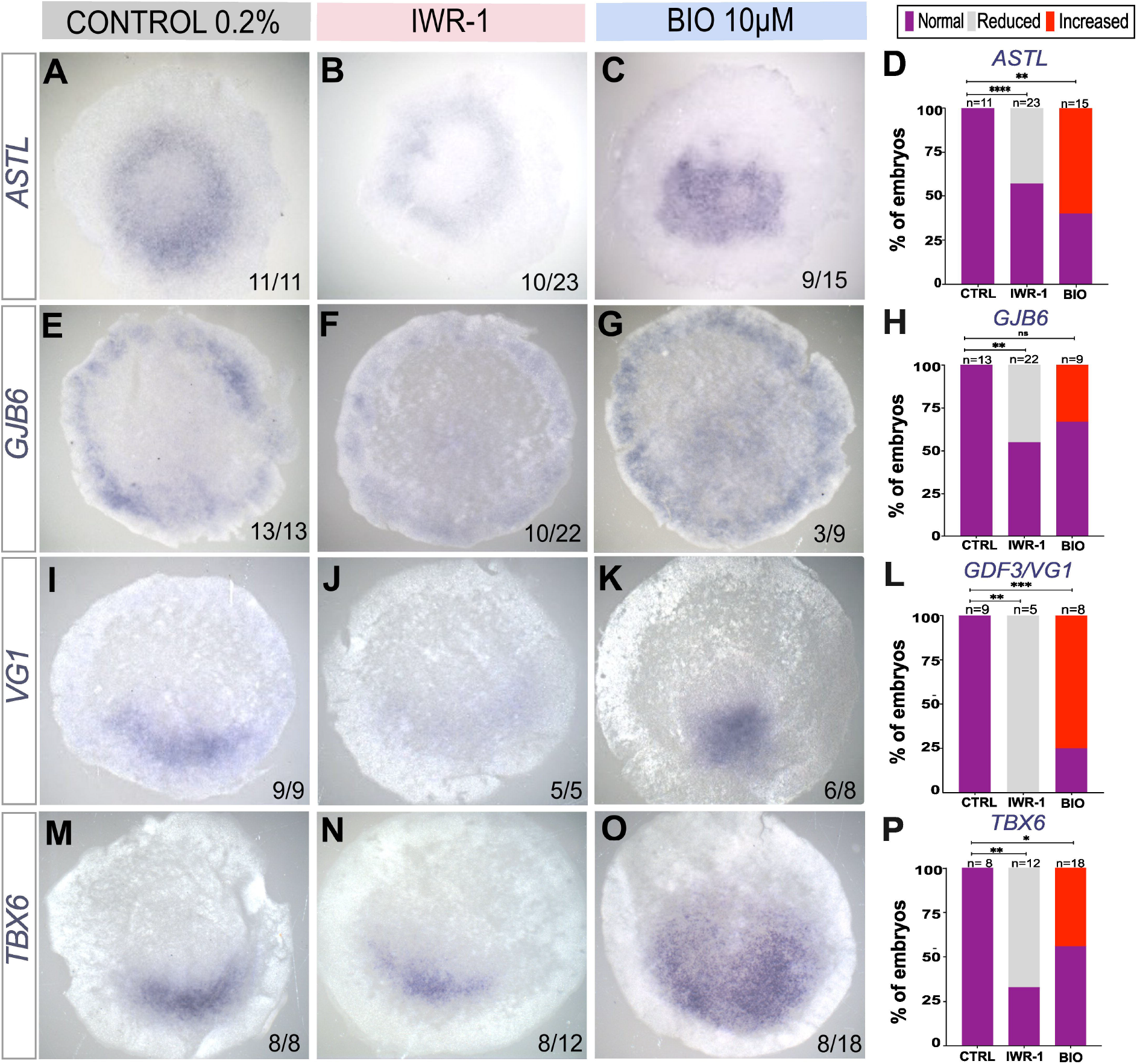
Wnt signalling regulates the MZ and posterior identity. **A**-**O**. In situ hybridisation showing expression of MZ related genes ASTL (A-C), GJB6 (E-G) and the posterior MZ marker Vg1/GDF3 (I-K) and TBX6 (M-O) in control (0.2% DMSO; A, E, I, M), IWR-1 (25μM) (B,F,J,N) and BIO (10μM) (C, G, K, O) treated EGK-XI embryos incubated for 6-hours. **D, H, L, P**. Stacked bar graphs showing percentage of embryos with normal, reduced, or increased expression compared to control of ASTL (D), GJB6 (H), VG1 (L) and TBX6 (P). P-value was calculated by Fisher’s exact (two side) test (ns) p-value > 0.05, (*) p-value <0.05, (**)p-value < 0.005, (***)p-value <0.0005, (****)p-value. <0.0001).

### The marginal zone is continuously and actively maintained by Wnt signalling

Is the continued presence of the AO required to maintain MZ properties in this region? To test this, we ablated the AO from EGK-XI embryos and incubated the AP+MZ explants for 7-hours (Supplementary Fig. S2A). In situ hybridisation revealed that all AP+MZ explants (9/9) lost *ASTL* expression (Supplementary Fig.S2C) compared to unoperated control embryos cultured for the same period (10/10) (Supplementary Fig.S2B). This indicates that signals from the AO are required to maintain MZ properties during normal development, after its initial formation.

To test whether Wnt signalling is responsible for maintenance of the MZ, we incubated whole, unoperated stage XI embryos in the Wnt antagonist IWR-1 for 6 hours, and analysed the embryos by *in situ* hybridisation. Expression of both *ASTL* and *GJB6* were inhibited: *ASTL* was reduced in 10/23 embryos (Fisher’s exact test; p-value<0.0001), and *GJB6* was reduced in 10/22 embryos (Fisher’s exact test; p-value=0.0052), respectively, compared to DMSO controls (Fig. 4 A, B, E, F). We also examined the expression of the posterior MZ markers *VG1* (*GDF3*) and *TBX6*: 5/5 IWR-1-treated embryos showed reduced expression of *VG1* (Fig. 4 J) (Fisher’s exact test; p-value=0.0005) and 8/12 embryos showed reduced expression of *TBX6* (Fig. 4 N) (Fisher’s exact test; p-value=0.0047) compared to the 0.2% DMSO control embryos, where 9/9 embryos had normal expression of *VG1* and 8/8 embryos were normal for *TBX6* expression (Fig. 4 I, M).

Treatment of whole EGK-XI embryos with BIO expanded the MZ into the AP, with broader *ASTL* (9/15 embryos) compared to the 0.2% DMSO incubated embryo where (11/11) embryos showed normal expression (Fisher’s exact test; p-value <0.0024) (Fig. 4 C). However, a lower proportion (3/9) of treated embryos had expansion of the *GJB6* expression domain (Fisher’s exact test; p-value < 0.0545) (Fig 4. G). Wnt stimulation also led to expansion of the posterior MZ into the AP, revealed by expansion of *VG1* (6/8 embryos) (Fisher’s exact test; p-value < 0.0023) and *TBX6* (8/18 embryos) (Fisher’s exact test; p-value < 0.038) compared to controls (Fig. 4 K, O, L, P). These findings suggest that Wnt activity is required for the maintenance of the MZ after its formation.

## Discussion

This study reveals that canonical Wnt-signalling can account for the recently reported induction of marginal zone (MZ) properties by the area opaca (AO) (Lee et al., 2022b). Inhibition of canonical Wnt signalling using IWR-1 prevents the AO from inducing MZ markers when recombined with the AP. Moreover, exposure of an isolated AP to the Wnt agonist BIO is sufficient to induce widespread expression of MZ markers. Furthermore, two findings suggest that continued signalling by the AO is required to maintain the MZ: first, ablation of the AO followed by culture of the AP+MZ regions alone causes MZ properties to be lost, and second, treatment of intact embryos (already containing a MZ) with IWR-1 result in the loss of MZ markers, suggests that the MZ is under active maintenance by Wnt signals until the start of gastrulation (primitive streak formation). These findings therefore suggest that Wnt signalling from the AO could be responsible for the initial induction of the MZ during normal development, which occurs at intrauterine stages of development. Consistent with these findings, we find Wnt ligands (*WNT-3A, -5A, -5B* and -*8C*) expressed in the AO, receptors strongly expressed in the AP. The MZ strongly expresses genes considered to be targets of Wnt signalling including *AXIN-2* and *LEF-1*, confirming earlier studies (Lee et al., 2022b, Roeser et al., 1999, Skromne and Stern, 2002). *WNT8C* is expressed most strongly and as a gradient, strongest posteriorly in the AO (Hume and Dodd, 1993, Skromne and Stern, 2001). Here we find that nuclear localisation of β-catenin, indicating where canonical signalling is active, is also graded in the MZ, with highest levels posteriorly, supporting the presence of a gradient.

A large number of studies in anamniote embryos has revealed that during very early stages of development, canonical Wnt activity is critical for defining the “dorsal” side of the embryo, thus defining the site where gastrulation will begin and therefore one of the axes of embryonic polarity (Larabell et al., 1997, Weaver et al., 2003, Salic et al., 2000, Miller et al., 1999, Dorsky et al., 2002, Lu et al., 2011). In these embryos, where the zygotic genome remains largely silent for the first 10 or so cell divisions, polarised expression is achieved by localization of maternal components like mRNAs or proteins – here, the dorsally-localized determinant is nuclear localization of β-catenin. In Xenopus, it is cortical rotation following fertilisation that generates the first dorsoventral difference, determining where the nuclear localization will take place (Larabell et al., 1997). The overlap between canonical Wnt activity and the transcription factor VegT (Tbx6) and the TGFβ/GDF-signalling component Vg1 defines the location of the future Nieuwkoop centre, responsible for induction of the organizer in adjacent cells (Brannon and Kimelman, 1996, Carnac et al., 1996, Fagotto et al., 1997).

In the chick embryo, an amniote, a region functionally equivalent to the Nieuwkoop centre has been located to the posterior MZ because it can induce the primitive streak including the organizer, without contributing any cells to the induced structures (Bachvarova et al., 1998), which are the defining hallmarks of the Nieuwkoop centre in amphibians (Harland and Gerhart, 1997, Nieuwkoop, 1969b, Nieuwkoop, 1969a, Nieuwkoop, 1973). Like the amphibian Nieuwkoop centre, the posterior MZ is a region of overlap of expression between a TGFβ/GDF component (*VG1/GDF3*) (Shah et al., 1997, Seleiro et al., 1996), the transcription factor TBX6 (Knezevic et al., 1997) and high canonical Wnt activity (this study). This is particularly interesting because unlike anamniotes, where this region of high Wnt activity is defined by maternal localization resulting from cortical rotation before the activation of zygotic gene expression (see above), in amniotes including the chick, the zygotic genome is activated very early, during the first few cleavage divisions. These results indicate that the “dorsal” localization of Wnt activity, by whatever mechanism, is a highly conserved feature of vertebrate development and essential for the establishment of embryo polarity that will position the site of gastrulation (dorsal blastopore in amphibians, primitive streak in amniotes).

Cooperation or synergy of TGFβ and canonical Wnt signalling seems to be even more highly conserved – even in the fly, several different developmental events involve both pathways (Bilder et al., 1998, Estella and Mann, 2008, Requena et al., 2017). An earlier study exploring the role of Vg1/GDF3 in primitive streak formation in the chick reported that when Vg1 is misexpressed anywhere in the MZ, it can induce an ectopic primitive streak in adjacent area pellucida epiblast; however, when Vg1-expressing cells are placed into the AP itself, no streak markers are induced (Skromne and Stern, 2001). Because of the conservation of Wnt/TGFβ interactions, the study then explored whether differences in Wnt activity could underlie this difference. Indeed, when Vg1/GDF3 is misexpressed in the MZ together with cells secreting Wnt antagonists like NFz8 this no longer induces an ectopic streak, whereas when cells expressing Wnt1 are co-transplanted with cells expressing Vg1/GDF3 into the AP, a region of Brachyury/TBXT expression (primitive streak marker) is induced next to the grafted cells (Skromne and Stern, 2001). Although that study was performed with transfected cells as the source of factors (and it is possible that other factors produced by the cells also contribute to the results), the paper proposed that interactions between Vg1 and Wnt are important in the induction of primitive streak formation in early chick development, as shown in Drosophila and other systems. Our present study opens a refinement of this interpretation: it is not just Wnt activity that is required for Vg1 to induce a primitive streak, but it may be that it is the MZ character induced by Wnt that is important. Thus, Wnt would define the ring of MZ properties all around the margin of the embryo, and the localization of Vg1 (and TBX6) would define its posterior part, where the inducing molecules are produced. These speculations are supported by the present findings that incubation of isolated AP fragments in the with Wnt agonist BIO induces expression of the marginal zone genes (*GJB6* and *ASTL*) and concomitant loss of the AP specific gap-junction marker *GJA1*.

We noted that the endogenous inhibitor of canonical Wnt signalling *DKK1* is expressed overlapping WnNT8C and its target *AXIN2* (Arendt and Nübler-Jung, 1999, Chapman et al., 2002, Foley et al., 2000, Lee et al., 2020, Skromne and Stern, 2002). Moreover, Wnt antagonists of the *sFRP (*secreted frizzled receptor protein) family are expressed in the AP (Supplementary Fig. S1E) (Lee et al., 2020, Skromne and Stern, 2002), and an additional Wnt modulator, *GPC-4* (glypican-4), which facilitates the diffusion of Wnt and may act as an inhibitor by binding the ligand (Ohkawara et al., 2003) is expressed in the hypoblast, just beneath the epiblast of the AP. These observations suggest that Wnt activity is more delicately controlled and positioned than merely by the localization of a Wnt ligand and its receptors.

In our previous study, the AO induced a MZ with uniformly posterior character, expressing Vg1 and TBX6 all the way around, even if the AO used was entirely anterior/lateral (Lee et al., 2022b). In the present study, we also see that isolated AO that has been uniformly treated with BIO contains a larger area of Vg1 expression.

## Materials and Methods

### Embryo harvest and New culture

Fertile Dekalb White hens’ eggs (Henry Stewart & Co., UK) were incubated at 38°C for 2 hours to obtain embryos at stages EGK X-XI. Embryos were explanted from the egg and manipulated in Pannett-Compton saline (Pannett and Compton, 1924), Phosphate Buffered Saline (PBS) or Tyrode’s solution (Tyrode, 1910) as previously described (Stern, 1993, Streit and Stern, 2008). They were then set up for modified New culture as previously described (New, 1955, Stern and Ireland, 1981) and incubated for the desired length of time as indicated in the text. Then they were fixed with 4% paraformaldehyde in Calcium/Magnesium-free PBS containing 2mM EGTA (pH 7.4) (PFA) overnight at 4°C. The next day, embryos are transferred to 100% methanol and stored for up to 3 days prior to in situ hybridisation.

### Pharmacological treatments

The Wnt agonist BIO (B1686; Sigma-Aldrich) and Wnt antagonist IWR-1 (I0161; Sigma-Aldrich) were stored as 5μl aliquots at 10mM and 25 mM respectively, in DMSO at □20°C. Two dilutions were used: 1:500 in PBS for pre-soaking (30 min at room temperature) and 1:1000 for New culture. Treatments were diluted in 1ml PBS then mixed with thin albumin to make up a final working concentration of 25μM for IWR-1 and 10 μM for BIO; 0.2% DMSO was used for controls.

### Embryo manipulations

Whole embryos were cultured for 6 hours to reach approximately stages EGK XII-XIII. For marginal zone removal experiments, the marginal zone was excised using a bent entomological pin (A1). The area opaca (after excision of about 45° arc from the posterior part) was then grafted onto the area pellucida and the two tissues sealed together by aspiration of as much liquid as possible using a fine micro-needle pulled from a 50 μl borosilicate capillary tube (Streit and Stern, 2008, Lee et al., 2022b). Operated embryos were cultured for 7 hours to reach stage EGK XIII-XIV. For area pellucida experiments, both the marginal zone and the area opaca were removed.

### Fixation and whole-mount in-situ hybridisation

Embryos were fixed in PFA overnight at 4°C. Whole mount in situ hybridisation was performed as previously described (Stern, 1998, Streit and Stern, 2001). The probes used were: *cVG1/GDF3* (Shah et al., 1997), *cBRA/TBXT* (Kispert et al., 1995), *ASTL* (ChEST817d16) (Lee et al., 2020), *GJB6* (ChEST89h10), *GJA1* (kind gift of Prof. Stephen Price), *WNT8C* (Hume and Dodd, 1993), *LEF-1* (Kengaku et al., 1998) and *TBX6* (Torlopp et al., 2014). Images of the stained embryos were taken using an Olympus SZH10 dissecting microscope with a QImaging Retiga 2000R camera using QCapture Pro software.

### β-catenin immunohistochemistry, confocal imaging and image analysis

Embryos at around stage EGK-XI were fixed in PFA for 1 hour, then washed with ice-cold methanol for 30 min. They were then rehydrated in a dilution series of methanol (75/50/25%) PBS containing 1% Triton-X100 (PBST), then washed for 15 min washes in PBST followed by 2-hour blocking in PBST containing 5% normal goat serum and 1% Thimerosal at room temperature. After blocking, embryos were incubated overnight at 4°C in mouse monoclonal antibody against β-catenin (1:200) (C7207; Sigma-Aldrich). The following day embryos were washed for 10-minute 3 times in PBST, followed by 3 longer 1-hour washes. Goat-anti mouse IgG Alexa-Fluor 594 (A11032; Invitrogen) was used as secondary antibody (1:500) in blocking buffer, with overnight incubation at 4°C. Embryos were then washed in PBST and embryos then counterstained for nuclei using 2.5 μg/ml DAPI. Next, embryos were mounted on a glass slide, ventral-side facing up using Vectashield (H-1000-10) mounting medium and overlain by a glass coverslip. Confocal imaging was done using a Leica SPE1 microscope. Images were processed using open-source FIJI (Image-J) software; segmentation was carried out using the DAPI channel of a 10-image z-stack, to create a region of interest (ROI) for the nuclei. Then, in the β-catenin channel, the pixel intensity of nuclear β-catenin was measured within the created ROI. A membrane ROI was created, using a 10-image z-stack of the β-catenin channel. The intensity of membrane β-catenin was measured using the ‘measure’ function in the ‘analyse particle’ menu. To normalize, we divided the pixel intensity of β-catenin in the membrane by the β-catenin intensity within the ROI of the nuclei.

### Graphs and Illustrations

All graphs and statistical analysis were done using GraphPad Prism version 9.5.1 (528). Line diagrams were generated using BioRender.com. For Supplementary Fig. 1A the illustrations were adapted from “Wnt signalling (active) and Wnt signalling (inactive)”, by BioRender.com (2024), retrieved from https://app.biorender.com/biorender-templates.

## Acknowledgements

This study was funded by a Wellcome Trust Investigator Award (107055/Z/15/Z) to C.D.S., which also supported H.C.L., H.-C.L. and N.M.M.O. H.C.L. was also supported by a fellowship from the Basic Science Research Program through the National Research Foundation of Korea (NRF) (2014R1A6A3A03053468). N.M.M.O. was also funded by UCL. Y.F. was a doctoral student in the Developmental and Stem Cell Biology Programme at UCL (Department of Cell and Developmental Biology) and was funded by a scholarship from the Custodian of The Two Holy Mosques’ External Scholarship Program from the Ministry of Education of Saudi Arabia (grant no. SE-12267).

## Figure Legends

**Supplementary Fig. S1. Wnt signalling modulation by Wnt agonist BIO and antagonist IWR-1. A**. Schematic diagram of ‘active’ and ‘inactive’ Wnt signalling, highlighting the presence of the degradation complex in the absence of the Wnt ligand, which ultimately degrades β-catenin, and when the Wnt is present β-catenin is stabilised and can translocate to the nucleus; the sites of action of BIO (as a GSK inhibitor, Wnt agonist) and IWR-1 (Tankyrase inhibitor, Wnt antagonist) are shown. **B-D**. In situ hybridisation of canonical Wnt targets after exposure to treatment for 7-hours. Control (0.2% DMSO, B), IWR-1 (C) and BIO (D). **E**. Stacked bar graph showing percentage of embryos with reduced, increased or normal expression the control sample (n=9) where 5 embryos are stage EGK-XII and 4 embryos stage HH2, (E) IWR-1 (25μM) n=8. P-value was determined using Fisher’s exact test; IWR-1 vs. Control p-value 0.0004 (***), BIO vs Control Fisher’s exact test p-value 0.0023 (**). (F-H) Confocal images of Fluorescent-IHC of β-catenin in the PAO of a z-stack maximum projection of EGK_XII treated embryos and control; DAPI=yellow, β-catenin=magenta. (F) A scattered bar plot showing the nuclear pixel intensity of β-catenin after 6-hours of incubation of stage EGK-XI embryos incubated with: 0.2% DMSO for the control, IWR-1 (25μM) and BIO (10μM) (n=2), each dot represents a segmented nucleus. Mann-Whitney U test P-value <0.0001; n=2. N: the number of embryos.

**Supplementary Fig. S2. The MZ is continuously maintained by the AO. A**. Schematic diagram of the experimental design. **B-C**. In situ hybridisation for MZ marker *ASTL* in normal, intact EGK-XI embryos (B) or EGK-XI embryos from which the AO had been removed and the MZ+AP fragment incubated (C).

**Supplementary Table S1. Ligand-receptor expression comparisons from CellChatDB analysis**.

## References

Arendt, D. & Nübler-Jung, K.1999. Rearranging gastrulation in the name of yolk: evolution of gastrulation in yolk-rich amniote eggs. Mech Dev, 81, 3–22.

Bachvarova, R. F., Skromne, I. & Stern, C. D.1998. Induction of primitive streak and Hensen’s node by the posterior marginal zone in the early chick embryo. Development, 125, 3521–34.

Bellairs, R., Bromham, D. R. & Wylie, C. C.1967. The influence of the area opaca on the development of the young chick embryo. J Embryol Exp Morphol, 17, 195–212.

Bilder, D., Graba, Y. & Scott, M. P.1998. Wnt and TGFbeta signals subdivide the AbdA Hox domain during Drosophila mesoderm patterning. Development, 125, 1781–90.

Brannon, M. & Kimelman, D.1996. Activation of Siamois by the Wnt pathway. Dev Biol, 180, 344–7.

Carnac, G., Kodjabachian, L., Gurdon, J. B. & Lemaire, P.1996. The homeobox gene Siamois is a target of the Wnt dorsalisation pathway and triggers organiser activity in the absence of mesoderm. Development, 122, 3055–65.

Chapman, S. C., Schubert, F. R., Schoenwolf, G. C. & Lumsden, A.2002. Analysis of spatial and temporal gene expression patterns in blastula and gastrula stage chick embryos. Dev Biol, 245, 187–99.

Dorsky, R. I., Sheldahl, L. C. & Moon, R. T.2002. A transgenic Lef1/beta-catenin-dependent reporter is expressed in spatially restricted domains throughout zebrafish development. Dev Biol, 241, 229–37.

Downie, J. R.1976. The mechanism of chick blastoderm expansion. J Embryol Exp Morphol, 35, 559–75.

Estella, C. & Mann, R. S.2008. Logic of Wg and Dpp induction of distal and medial fates in the Drosophila leg. Development, 135, 627–36.

Eyal-Giladi, H. & Khaner, O.1989. The chick’s marginal zone and primitive streak formation. II. Quantification of the marginal zone’s potencies--temporal and spatial aspects. Dev Biol, 134, 215–21.

Eyal-Giladi, H. & Kochav, S.1976. From cleavage to primitive streak formation: a complementary normal table and a new look at the first stages of the development of the chick. I. General morphology. Dev Biol, 49, 321–37.

Fagotto, F., Guger, K. & Gumbiner, B. M.1997. Induction of the primary dorsalizing center in Xenopus by the Wnt/GSK/beta-catenin signaling pathway, but not by Vg1, Activin or Noggin. Development, 124, 453–60.

Foley, A. C., Skromne, I. & Stern, C. D.2000. Reconciling different models of forebrain induction and patterning: a dual role for the hypoblast. Development, 127, 3839–54.

Funayama, N., Fagotto, F., Mccrea, P. & Gumbiner, B. M.1995. Embryonic axis induction by the armadillo repeat domain of beta-catenin: evidence for intracellular signaling. J Cell Biol, 128, 959–68.

Guger, K. A. & Gumbiner, B. M.1995. beta-Catenin has Wnt-like activity and mimics the Nieuwkoop signaling center in Xenopus dorsal-ventral patterning. Dev Biol, 172, 115–25.

Hamburger, V. & Hamilton, H. L.1951. A series of normal stages in the development of the chick embryo. J Morphol, 88, 49–92.

Harland, R. & Gerhart, J.1997. Formation and function of Spemann’s organizer. Annu Rev Cell Dev Biol, 13, 611–67.

Hatada, Y. & Stern, C. D.1994. A fate map of the epiblast of the early chick embryo. Development, 120, 2879–89.

He, X., Saint-Jeannet, J. P., Woodgett, J. R., Varmus, H. E. & Dawid, I. B.1995. Glycogen synthase kinase-3 and dorsoventral patterning in Xenopus embryos. Nature, 374, 617–22.

Hume, C. R. & Dodd, J.1993. Cwnt-8C: a novel Wnt gene with a potential role in primitive streak formation and hindbrain organization. Development, 119, 1147–60.

Jin, S., Guerrero-Juarez, C. F., Zhang, L., Chang, I., Ramos, R., Kuan, C.-H., Myung, P., Plikus, M. V. & Nie, Q.2021. Inference and analysis of cell-cell communication using CellChat. Nature Communications, 12, 1088.

Kengaku, M., Capdevila, J., Rodriguez-Esteban, C., De La Pena, J., Johnson, R. L., Izpisua Belmonte, J. C. & Tabin, C. J.1998. Distinct WNT pathways regulating AER formation and dorsoventral polarity in the chick limb bud. Science, 280, 1274–7.

Kispert, A., Ortner, H., Cooke, J. & Herrmann, B. G.1995. The chick Brachyury gene: developmental expression pattern and response to axial induction by localized activin. Dev Biol, 168, 406–15.

Knezevic, V., De Santo, R. & Mackem, S.1997. Two novel chick T-box genes related to mouse Brachyury are expressed in different, non-overlapping mesodermal domains during gastrulation. Development, 124, 411–9.

Larabell, C. A., Torres, M., Rowning, B. A., Yost, C., Miller, J. R., Wu, M., Kimelman, D. & Moon, R. T.1997. Establishment of the dorso-ventral axis in Xenopus embryos is presaged by early asymmetries in beta-catenin that are modulated by the Wnt signaling pathway. J Cell Biol, 136, 1123–36.

Lee, H. C., Fadaili, Y. & Stern, C. D. 2022a. Molecular characteristics of the edge cells responsible for expansion of the chick embryo on the vitelline membrane. Open Biol, 12, 220147.

Lee, H. C., Fadaili, Y. & Stern, C. D. 2024a. Development and functions of the area opaca of the chick embryo. (submitted).

Lee, H. C., Hastings, C. & Stern, C. D. 2022b. The extra-embryonic area opaca plays a role in positioning the primitive streak of the early chick embryo. Development, 149.

Lee, H. C., Lu, H. C., Turmaine, M., Oliveira, N. M. M., Yang, Y., De Almeida, I. & Stern, C. D.2020. Molecular anatomy of the pre-primitive-streak chick embryo. Open Biol, 10, 190299.

Lee, H. C., Oliveira, N. M. M., Hastings, C., Baillie-Benson, P., Moverley, A. A., Lu, H. C., Zheng, Y., Wilby, E. L., Weil, T. T., Page, K. M., Fu, J., Moris, N. & Stern, C. D. 2024b. Regulation of long-range BMP gradients and embryonic polarity by propagation of local calcium-firing activity. Nat Commun, 15, 1463.

Lu, F. I., Thisse, C. & Thisse, B.2011. Identification and mechanism of regulation of the zebrafish dorsal determinant. Proc Natl Acad Sci U S A, 108, 15876–80.

Martins-Neves, S. R., Paiva-Oliveira, D. I., Fontes-Ribeiro, C., Bovee, J., Cleton-Jansen, A. M. & Gomes, C. M. F.2018. IWR-1, a tankyrase inhibitor, attenuates Wnt/beta-catenin signaling in cancer stem-like cells and inhibits in vivo the growth of a subcutaneous human osteosarcoma xenograft. Cancer Lett, 414, 1–15.

Meijer, L., Skaltsounis, A. L., Magiatis, P., Polychronopoulos, P., Knockaert, M., Leost, M., Ryan, X. P., Vonica, C. A., Brivanlou, A., Dajani, R., Crovace, C., Tarricone, C., Musacchio, A., Roe, S. M., Pearl, L. & Greengard, P.2003. GSK-3-selective inhibitors derived from Tyrian purple indirubins. Chem Biol, 10, 1255–66.

Miller, J. R., Rowning, B. A., Larabell, C. A., Yang-Snyder, J. A., Bates, R. L. & Moon, R. T.1999. Establishment of the dorsal-ventral axis in Xenopus embryos coincides with the dorsal enrichment of dishevelled that is dependent on cortical rotation. J Cell Biol, 146, 427–37.

Narwal, M., Venkannagari, H. & Lehtio, L.2012. Structural basis of selective inhibition of human tankyrases. J Med Chem, 55, 1360–7.

New, D. A. T.1955. A new technique for the cultivation of the chick embryo in vitro. J. Embryol. exp. Morph., 3, 326–331.

Nieuwkoop, P. D. 1969a. The formation of mesoderm in urodelean amphibians I. Induction by the endoderm. W Roux Arch EntwMech Org, 162, 341–373.

Nieuwkoop, P. D. 1969b. The formation of mesoderm in Urodelean amphibians. II. The origin of the dorso-ventral polarity of the mesoderm. Wilh Roux’ Arch EntwMech Organ, 163, 298–315.

Nieuwkoop, P. D.1973. The organization center of the amphibian embryo: its origin, spatial organization, and morphogenetic action. Adv Morphog, 10, 1–39.

Ohkawara, B., Yamamoto, T. S., Tada, M. & Ueno, N.2003. Role of glypican 4 in the regulation of convergent extension movements during gastrulation in Xenopus laevis. Development, 130, 2129–38.

Pannett, C. & Compton, A.1924. THE CULTIVATION OF TISSUES IN SALINE EMBRYONIC JUICE. The Lancet, 203, 381–384.

Requena, D., Alvarez, J. A., Gabilondo, H., Loker, R., Mann, R. S. & Estella, C.2017. Origins and Specification of the Drosophila Wing. Curr Biol, 27, 3826–3836 e5.

Roeser, T., Stein, S. & Kessel, M.1999. Nuclear beta-catenin and the development of bilateral symmetry in normal and LiCl-exposed chick embryos. Development, 126, 2955–65.

Salic, A., Lee, E., Mayer, L. & Kirschner, M. W.2000. Control of beta-catenin stability: reconstitution of the cytoplasmic steps of the wnt pathway in Xenopus egg extracts. Mol Cell, 5, 523–32.

Schmidt, M., Patterson, M., Farrell, E. & Munsterberg, A.2004. Dynamic expression of Lef/Tcf family members and beta-catenin during chick gastrulation, neurulation, and early limb development. Dev Dyn, 229, 703–7.

Schneider, S., Steinbeisser, H., Warga, R. M. & Hausen, P.1996. Beta-catenin translocation into nuclei demarcates the dorsalizing centers in frog and fish embryos. Mech Dev, 57, 191–8.

Seleiro, E. A., Connolly, D. J. & Cooke, J.1996. Early developmental expression and experimental axis determination by the chicken Vg1 gene. Curr Biol, 6, 1476–86.

Shah, S. B., Skromne, I., Hume, C. R., Kessler, D. S., Lee, K. J., Stern, C. D. & Dodd, J.1997. Misexpression of chick Vg1 in the marginal zone induces primitive streak formation. Development, 124, 5127–38.

Skromne, I. & Stern, C. D.2001. Interactions between Wnt and Vg1 signalling pathways initiate primitive streak formation in the chick embryo. Development, 128, 2915–27.

Skromne, I. & Stern, C. D.2002. A hierarchy of gene expression accompanying induction of the primitive streak by Vg1 in the chick embryo. Mech Dev, 114, 115–8.

Stern, C. D.1993. Avian embryos. In: Stern, C. D. & Holland, P. W. H. (eds.) Essential Developmental Biology: A Practical Approach. Oxford: IRL Press/Oxford University Press.

Stern, C. D.1998. Detection of multiple gene products simultaneously by in situ hybridization and immunohistochemistry in whole mounts of avian embryos. Curr Top Dev Biol, 36, 223–43.

Stern, C. D. & Ireland, G. W.1981. An integrated experimental study of endoderm formation in avian embryos. Anat Embryol (Berl), 163, 245–63.

Streit, A. & Stern, C. D.2001. Combined whole-mount in situ hybridization and immunohistochemistry in avian embryos. Methods, 23, 339–44.

Streit, A. & Stern, C. D.2008. Operations on primitive streak stage avian embryos. Methods Cell Biol, 87, 3–17.

Torlopp, A., Khan, M. A., Oliveira, N. M., Lekk, I., Soto-Jimenez, L. M., Sosinsky, A. & Stern, C. D.2014. The transcription factor Pitx2 positions the embryonic axis and regulates twinning. Elife, 3, e03743.

Tyrode, M. V.1910. The Mode of Action of Some Regulative Salts. 17, 205–209.

Weaver, C., Farr, G. H., 3RD, Pan, W., Rowning, B. A., Wang, J., Mao, J., Wu, D., Li, L., Larabell, C. A. & Kimelman, D.2003. GBP binds kinesin light chain and translocates during cortical rotation in Xenopus eggs. Development, 130, 5425–36.

Yost, C., Farr, G. H., 3RD, Pierce, S. B., Ferkey, D. M., Chen, M. M. & Kimelman, D.1998. GBP, an inhibitor of GSK-3, is implicated in Xenopus development and oncogenesis. Cell, 93, 1031–41.

Yost, C., Torres, M., Miller, J. R., Huang, E., Kimelman, D. & Moon, R. T.1996. The axis-inducing activity, stability, and subcellular distribution of beta-catenin is regulated in Xenopus embryos by glycogen synthase kinase 3. Genes Dev, 10, 1443–54.

