## Supplementary figures and images for "Canonical Wnt signalling from the Area Opaca induces and maintains the Marginal Zone in pre-primitive-streak stage chick embryos"

### Supplementary Figure S1

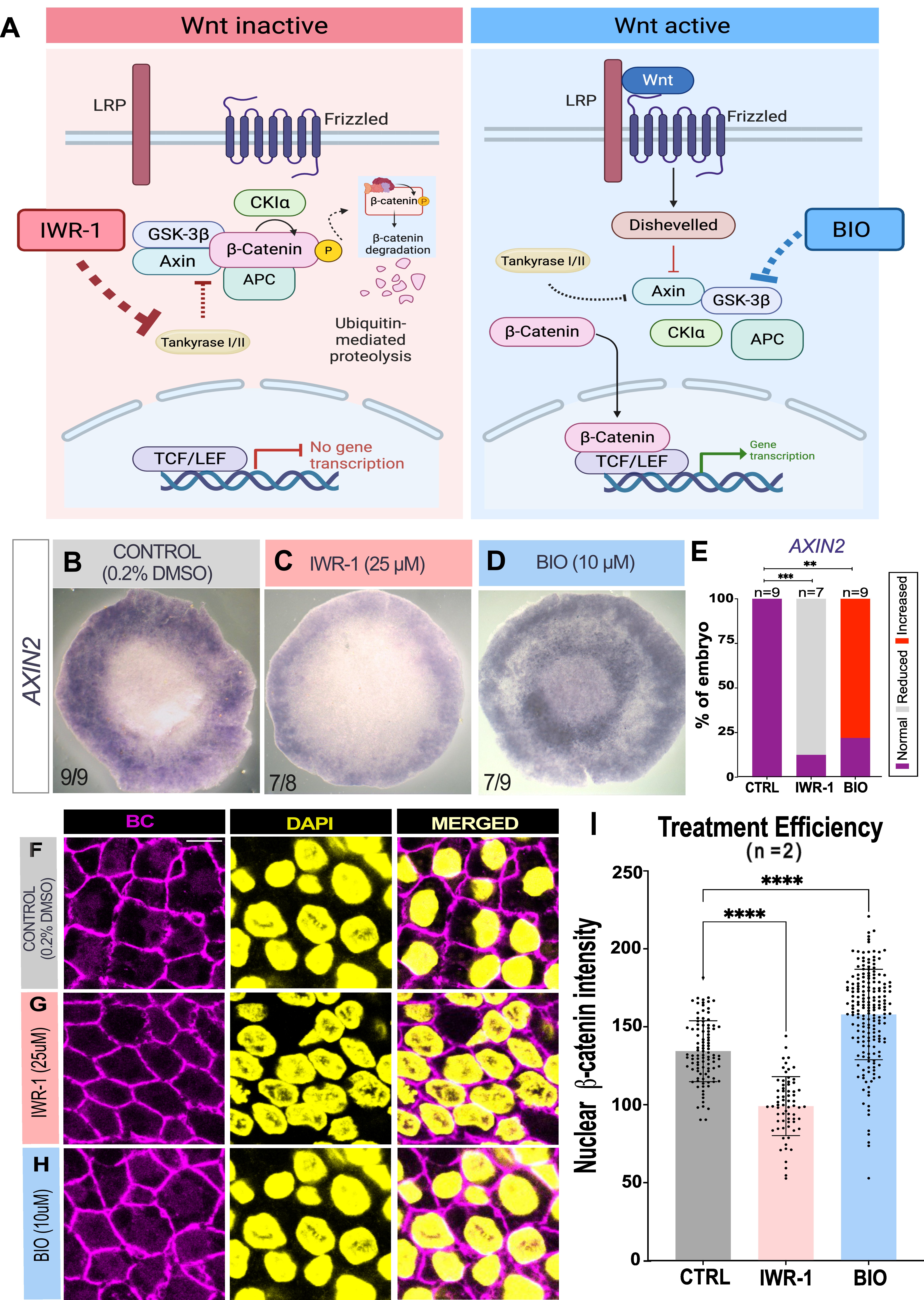

### Supplementary Figure S2

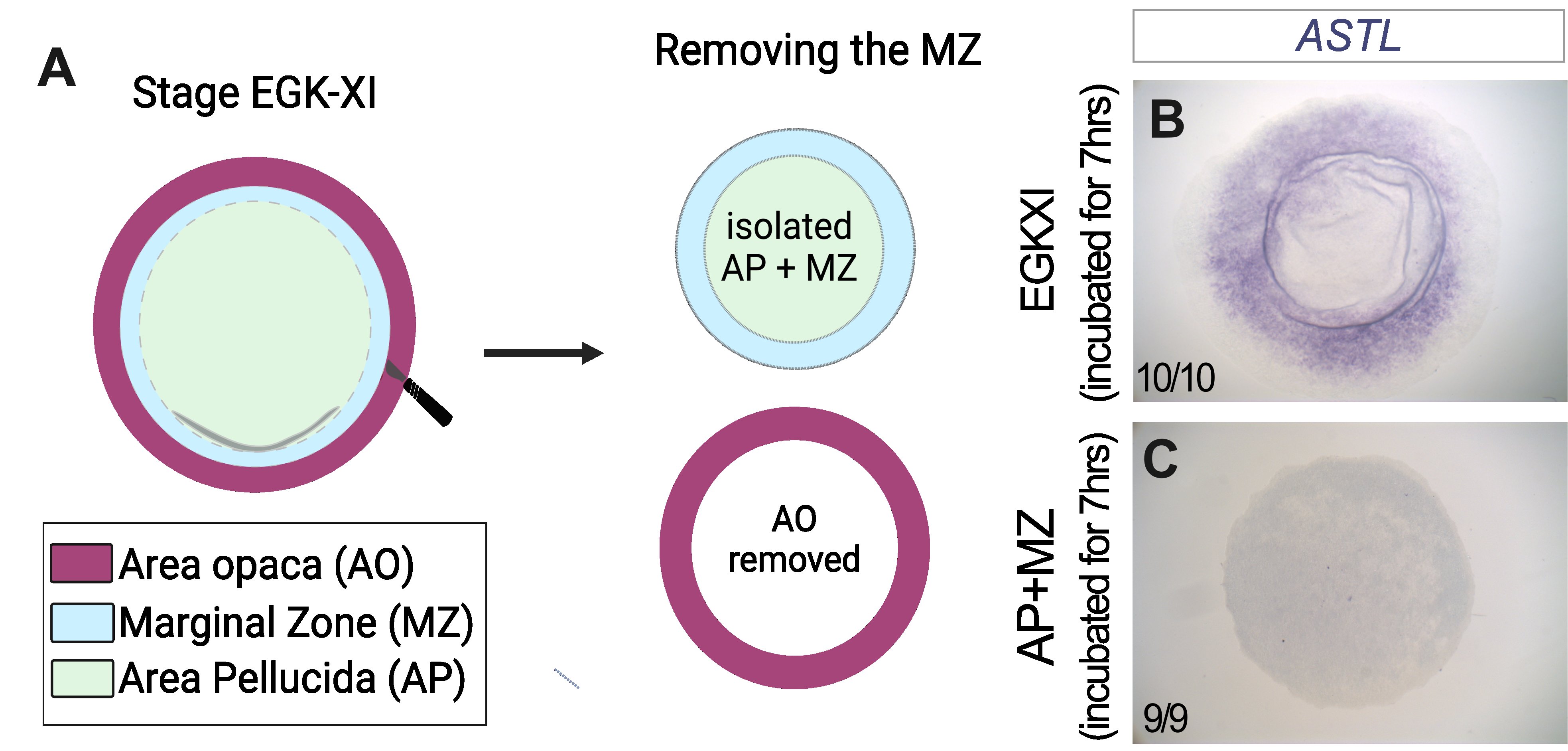
